# Outcome-selective reinstatement is predominantly context-independent, and associated with c-Fos activation in the posterior dorsomedial striatum

**DOI:** 10.1101/2021.07.28.454246

**Authors:** Arvie R. Abiero, Zaid Ali, Bryce Vissel, Laura A. Bradfield

**Affiliations:** Centre for Neuroscience and Regenerative Medicine, School of Life Sciences, Faculty of Science, University of Technology Sydney, New South Wales, 2007, Australia; St. Vincent’s Centre for Applied Medical Research, St. Vincent’s Health Network, Sydney, New South Wales, 2010, Australia

**Author notes:** **Co-Senior authorship** (Bryce Vissel, Laura A. Bradfield). **Corresponding Author** Dr. Laura A. Bradfield, Centre for Neuroscience and Regenerative Medicine, School of Life Sciences, Faculty of Science, University of Technology Sydney (St. Vincent’s Campus), 405 Liverpool Street, Darlinghurst, NSW 2010.

**Keywords:** Contextual modulation, dorsomedial striatum, outcome-selective reinstatement, substance use disorder, c-Fos activation

## Abstract

Research from human and animal studies has found that responding that has been successfully reduced following treatment can return upon exposure to certain contexts. An individual in recovery from alcohol use disorder, for example, might relapse to drinking upon visiting their favourite bar. However, most of these data have been derived from experiments involving a single (active) response, and the context-dependence of returned responding in situations involving choice between multiple actions and outcomes is less well-understood. We thus investigated how outcome-selective reinstatement – a procedure involving choice between multiple actions – was affected by altering the physical context in rats. In Experiment 1, rats were trained over 6 days to press a left lever for one food outcome (pellets or sucrose) and a right lever for the other outcome. Then, rats received an extinction session in either the same context (A) as lever press training, or in a different context (B). Rats were tested immediately (5 minutes) after extinction in Context A or B such that there were four groups in total: AAA, ABB, ABA, and AAB. Reinstatement testing consisted of one food outcome being delivered ‘freely’ (i.e. unearned by lever pressing and unsignalled by cues) to the food magazine every 4 minutes in the following order: Sucrose, Pellet, Pellet, Sucrose. Selective reinstatement was considered intact if pellet delivery increased pressing selectively on the pellet lever, and sucrose delivery selectively increased pressing on the sucrose lever. This result (Reinstated > Nonreinstated) was observed for rats in group AAA and ABB, but not rats in groups ABA and AAB. Experiment 2 was conducted identically, except that rats received two extinction sessions over two days and tested one day later. This time, all groups demonstrated intact outcome-selective reinstatement regardless of context. Analysis of c-Fos expression in several brain regions revealed that only c-Fos expression in the posterior dorsomedial striatum (pDMS) was related to intact reinstatement performance. Overall, these results suggest that outcome-selective reinstatement is predominantly context-independent, and that intact reinstatement is related to neuronal activity in the pDMS.

**Highlights:**

- Outcome-selective reinstatement is predominantly context-independent
- Outcome-selective reinstatement is entirely context-independent after multiple extinction sessions
- Outcome-selective reinstatement increases c-Fos expression in dorsomedial striatum
- c-Fos expression in orbitofrontal cortex and dorsal hippocampus is unaffected by selective reinstatement.

## 1. Introduction

The increase in responding that is observed after a period of treatment or intervention, such as a return to drug seeking following a treatment in substance use disorder (SUD), is commonly modelled in animals through procedures such as renewal and reinstatement. Such models are essential because they allow researchers to gain a deeper understanding of the behavioural and brain mechanism of relapse using techniques that are not viable for use in human studies. In these procedures, the animal is typically first trained to administer a drug, food, or other rewarding substance, which is earned by performing a response such as lever pressing. Treatment is then modelled via a process known as ‘extinction’ during which responding no longer earns the desired outcome until it subsequently declines. Reinstatement is observed if the animal is later subject to an unsignalled, unearned delivery of the outcome, causing responding to re-emerge (de Wit & Stewart, 1981; Stretch & Gerber, 1973). Renewal (or context-induced reinstatement), on the other hand, is the increase in responding that is observed if the animal is placed into a context other than that in which extinction took place (Bouton et al., 2011; Delamater, 1997).

Although renewal and reinstatement are useful models, they do not fully capture the richness of the relapse environment. This is because the majority of studies using these models require animals to make a single, active response for a single outcome. In the real world, however, an individual performing a single response for a single outcome in a uniform environment is rare. Rather, as noted in a recent review (Vandaele & Ahmed, 2021), people are far more likely to be faced with scenarios in which they can choose between multiple actions that have multiple outcomes. A person who both drinks and smokes, for example, will have many context and response-bound associations with both behaviours (e.g. at bars, the balcony at home, garden at work etc). Thus, it is the aim of this study to begin to capture some of this complexity at a preclinical level, by determining how selective reinstatement in a choice-based procedure is influenced by context. Specifically, we ask whether increases in multiple responses for multiple outcomes, rather than selective responding for a single outcome, is more likely after a change in physical context. For example, would a person who has been through treatment, and thus abstinent from both alcohol and cigarettes, but then relapses on one of these drugs in their local bar also be more likely to relapse on the other? And are they more likely to relapse on both in this context than if they were in another, more neutral context? In order to answer this question whilst avoiding the potentially confounding effects that drugs have on the physical and/or mental state of the animals, we used food rather than drug outcomes for the current study.

The phenomenon of renewal was first demonstrated in Pavlovian (i.e. stimulus-outcome) conditioning (Bouton & Bolles, 1979), and later replicated in instrumental (i.e. action-outcome) conditioning (e.g. Bouton et al., 2011). There are several ways in which renewal can occur, but ABA renewal is the most common and most robust. In ABA renewal, a stimulus or response is initially paired with an outcome in Context A, then extinguished in a different context, Context B, after which the animal is returned to Context A where responding tends to increase again. This increase is apparent when their responding is compared to animals who were trained, extinguished, and tested all in the same context (AAA), or trained in one context, but extinguished and tested in another (ABB). ABA renewal is thought to occur because it reduces the ambiguity that results from the extinction procedure, after which the cue or action has been both paired with the outcome and with nothing. If the cue/action has been consistently paired with the outcome in one context (A) but not another (B), however, this contextual information can reduce ambiguity such that when the cue/action is presented again in Context A responding is increased in line with the expectation of reinforcement (e.g. Bouton et al., 1994; 2011; 2021). Other forms of renewal have also been demonstrated in instrumental conditioning paradigms, such as AAB renewal which is observed when animals are trained and extinguished in the same context (A) and then tested in a different context (B) where responding returns.

Reinstatement, like renewal, is thought to be a contextually-mediated phenomenon (Delamater, 1997). Specifically, when the outcome is delivered on test in a manner that is unsignalled – i.e. unpaired with the response – this is thought to imply that the outcome is once again available within that context which increases the propensity to respond. Consistent with this idea, reinstatement is reduced if the outcome is presented in a context different to that of initial training or test (Baker et al., 1991). However, an alternative account has been proposed in which the outcome itself could provide a local context of reinforcement. For instance, when rats learn to press a lever for pellets, they will often retrieve and consume a pellet then shortly afterwards perform their next lever press. This is thought to lead to the formation of pellet-lever press (i.e. outcome-response) associations as well as the more traditionally considered lever press-pellet (i.e. response-outcome) associations. When the pellet is later delivered on test in the absence of a preceding lever press, it could activate these outcome-response associations in such a way the outcome acts as a stimulus that drives, or sets the occasion for, the response regardless of the physical context (Bouton et al., 2021).

Selective reinstatement in a two action, two outcome paradigm has been explicitly demonstrated to rely on outcome-response associations. Ostlund and Balleine (2007) first trained rats to press a left lever for pellets and a right lever for sucrose (or vice versa, counterbalanced) then gave rats an extinction session during which neither lever earned any outcome. Immediately after extinction, rats received two unsignalled presentations of each outcome, separated by several minutes during which lever presses were recorded but did not earn any outcomes. Reinstatement during these post-outcome delivery periods was found to be selective, because sucrose delivery selectively elicited responding on the sucrose lever, and pellet delivery elicited presses on the pellet lever. Ostlund and Balleine further demonstrated that shifting the current incentive value of the outcome (i.e. devaluing it through specific satiety) did not alter the selectivity of reinstatement, suggesting that the outcome was not functioning as a goal of lever pressing in this paradigm. In a final experiment, Ostlund and Balleine trained each outcome to explicitly function as both a stimulus for, and a goal of, lever pressing. For animals in the congruent group, the contingencies were: pellet: (left lever→pellet), sucrose: (right lever→sucrose), or vice versa, counterbalanced. This group should reinstate on the same lever regardless of whether the outcome functioned as a goal or stimulus. For the incongruent group, rats were trained so that pellet: (left lever→sucrose), sucrose: (right lever→pellet) (or vice versa, counterbalanced). In this group, if the outcome functioned as a stimulus during reinstatement then animals should have reinstated responding on the lever it preceded (in the above example this would be pellet→ left lever, sucrose→right lever), whereas if it were functioning as a goal, this group should reinstate responding on the lever press that earned it during training (i.e. pellet→right lever, sucrose→left lever). When tested, the incongruent group clearly reinstated on the lever that the outcome had preceded, demonstrating that the outcome was functioning as a stimulus, and suggesting that outcome-response associations rather than response-outcome associations underlie the selective reinstatement effect (Ostlund & Balline, 2007).

Because selective reinstatement depends on outcome-response associations in which the outcome functions as a stimulus, this suggests that it is more akin to habitual responding which also relies on stimulus-response associations than goal-directed actions that rely on response-outcome associations (Balleine & Dickinson, 1998). This is notable for the current study, because habits have been shown to depend on the context in which they are learned (Bouton et al., 2021; Bouton et al., 2011) whereas goal-directed actions that have been demonstrated to be relatively context-independent (Bradfield et al., 2020; Thrailkill & Bouton, 2015). Thus, if context-specificity is a general property of stimulus-response associations we might expect outcome-selective reinstatement to also be context-specific. Alternatively, it is possible that outcome-response associations do not function in the same way as other stimulus-response associations, and are in fact independent of their learning context, consistent with the suggestion by Bouton et. al., (2021) that it is the outcome itself that provides the ‘context’ for reinstatement.

The current study will test these opposing predictions. We will employ the same outcome-selective reinstatement paradigm used by Ostlund & Balleine (2007), except that the physical contexts will be altered during extinction and test. Specifically, lever press training will take place in Context A for all groups, whereas extinction and testing will take place in either Context A or B, yielding four groups in total: groups AAA, ABB, ABA, and AAB. Intact selective reinstatement will be indicated by greater responding on the reinstated lever than the nonreinstated lever (Reinstated > Nonreinstated), and impaired selective reinstatement will be indicated by equivalent responding on both levers (Reinstated = Nonreinstated). If selective reinstatement is context-dependent, it should be intact for group AAA, for whom the context does not vary, and should be impaired, for group ABB for whom training and testing occurs in different contexts. If it is context-independent, however, it should be intact in both groups. We additionally included ‘renewal’ groups AAB and ABA, for whom the contexts were switched between extinction and test. Based on previous demonstrations of AAB and ABA renewal in instrumental conditioning (Bouton et al., 2011), we expected that responding would increase for these groups when placed into the test context in a manner that should interfere with the selective reinstatement effect. That is, because reinstatement depends on the reduction in responding that occurs as a result of extinction learning, then removing the influence of extinction learning via renewal should increase responding and preclude its observation. Thus, selective reinstatement was expected to be impaired for these groups. Experiments 1 and 2 were conducted identically, except that rats were tested immediately after extinction in Experiment 1, in line with the procedures employed by Ostlund and Balleine (2007), whereas rats were tested using more traditional extinction parameters (i.e. two extinction sessions and tested one day later) in Experiment 2, to determine the generality of the observed effects.

Finally, because relatively little is known about the neural mechanisms underlying outcome-selective reinstatement, we aimed to identify its potential neural correlates by examining expression of the immediate early gene and activity marker c-Fos in various brain regions. One brain region that selective reinstatement has been demonstrated to depend upon is the posterior dorsomedial striatum (pDMS) (Yin et al., 2005) and one region it does not depend upon is the medial orbitofrontal cortex (mOFC) (Bradfield et al., 2015; Bradfield et al., 2018). Thus, we included both regions in our analysis with the expectation that c-Fos expression would reflect performance in the pDMS but not the mOFC. We additionally conducted exploratory analyses of c-Fos expression in the lateral OFC (lOFC), given its central role in a varied number of reinforcement learning paradigms, as well as the dorsal hippocampus (DH) given its central role in contextual representation and learning.

## 2. Experiment 1: When tested immediately flowing extinction, outcome-selective reinstatement is context-independent, but extinction learning is not

The aim of Experiment 1 was to determine whether outcome-selective reinstatement is context dependent, and to investigate the potential neural mechanisms of this effect. The design for this experiment is in Figure 1. All rats were trained to lever press in Context A, the identity of which was counterbalanced. Half of the rats in each group were trained to press a left lever for pellets and a right lever for a sucrose solution, and the other half were trained on the opposite contingencies. All rats then received 30 minutes of extinction during which both levers were extended but no outcomes delivered. For groups AAA and AAB, this extinction session occurred in the same context as lever press training (Context A) whereas for groups ABB and ABA it occurred in a context that had different wallpaper, floor texture, and odour (Context B). Following extinction, rats were briefly put back into their home cage for 5 minutes to allow the experimenter to change the context adorning the operant chamber, then immediately transferred back into the operant chamber for testing.

**Figure 1.**
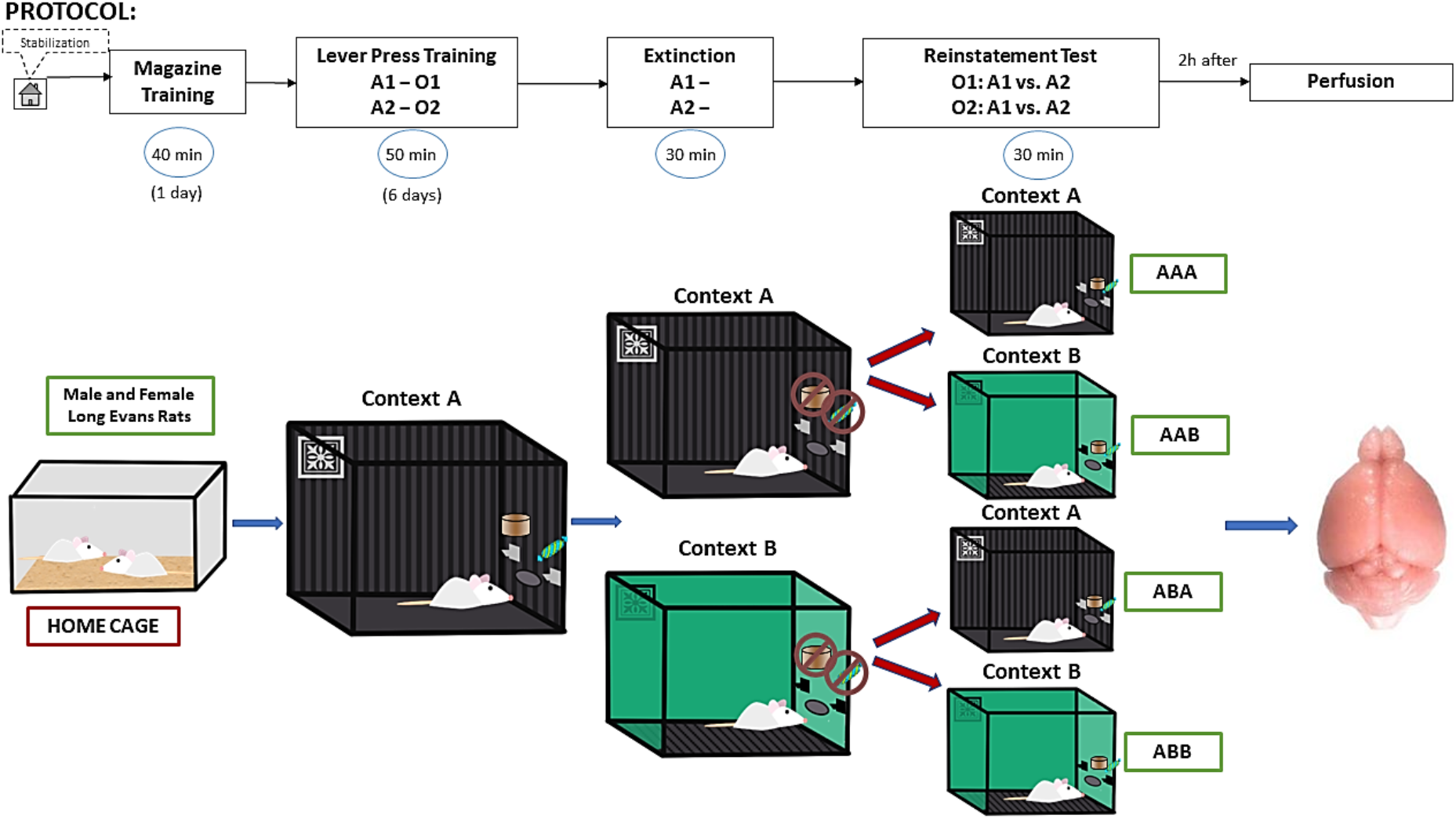
Representation of the experimental design. Briefly, animals were first trained to press left lever-pellets, right lever-sucrose, or vice versa (counterbalanced) in Context A. Next, lever pressing was extinguished in either the same context (A) or different context (B). Subsequently, reinstatement responding was assessed in context A or B, yielding 4 groups in total: AAA, ABB, ABA, and AAB.

Testing occurred in either Context A (groups AAA and ABA) or B (groups ABB and AAB). No outcomes were delivered in the first 3 minutes of the test session to allow us to measure whether there were any increases in responding as a result of the alterations in context alone – i.e. to measure renewal prior to reinstatement. Following this 3-minute period, rats received sucrose, pellet, pellet, sucrose, presentations in that order, each separated by 4 minutes of extinction. Baseline lever pressing was recorded for the 2 minutes prior to each outcome delivery, and selective reinstatement was measured as the number of lever presses on the reinstated lever compared to presses on the nonreinstated lever in the 2 minutes post-outcome delivery. We expected that selective reinstatement (Reinstated > Nonreinstated) would be intact in group AAA, but impaired (Reinstated = Nonreinstated) in all other groups.

Two hours after the start of the reinstatement test animals were perfused with 4% paraformaldehyde, their brains removed, and sections from the mOFC, lOFC, pDMS, and DH were extracted and immunostained for expressed of the immediate early gene and neuronal activity marker c-Fos. It was expected that c-Fos expression in the pDMS would be higher in groups expressing intact outcome-selective reinstatement than those for which it was impaired, whereas it was expected to be equivalent in the mOFC in all groups. Because the role of lOFC and DH in selective reinstatement is unknown, these analyses were exploratory, but it was expected that c-Fos expression in DH would differ in groups that experienced a context change on test day (groups AAB and ABA) relative to those that did not (groups AAA and ABB).

## 3. Materials and Methods – Experiment 1

### 3.1. Animals

A total of 20 male and 20 female Long-Evans rats were used for Experiment 1 (n = 11 (AAA), n = 10 (ABB), n = 10 (ABA), and n = 9 (AAB), N = 40). Male and female rats were assigned evenly to each group. The slightly uneven group numbers in AAA and AAB in Experiment 1 were due to one animal being incorrectly assigned to group AAA instead of AAB on test. All animals were purchased from the Australian Research Centre, Perth, Australia, and were housed in groups of 2-4 in transparent amber plastic boxes located in a temperature- and humidity-controlled room with a 12-h light/dark (07:00–19:00 h light) schedule. Experiments were conducted during the light cycle. Rats were aged between 10-15 weeks and weighed between 190-250 g (female) or 290-380 g (male) at the beginning of the experiment. Before the experiments, all animals were habituated to the housing area for a week, during which they had free access to food and water. During behavioural training however, animals were maintained at ~85% of their free-feeding body weight by limiting their food intake to 8-12g of their maintenance diet per day. All procedures were approved by the Ethics Committees of the Garvan Institute of Medical Research, Sydney.

### 3.2. Apparatus

All behavioural procedures took place in six identical sound attenuating operant chambers (Med Associates, Inc.,) that were enclosed in sound- and light-attenuating cubicles. Each chamber was equipped with a recessed food magazine, located at the base of one end wall, through which 20% sucrose solution (0.2 ml) and food pellets (45 mg; Bio-Serve, Frenchtown, NJ) could be delivered using a syringe pump and pellet dispenser, respectively. Pellet and sucrose outcomes were delivered to the same food magazine, in separate compartments to prevent pellets becoming wet. An infrared light situated at the magazine opening was used to detect head entries. Illumination was provided by a 3-W, 24-V house situated at the top-centred on the left end wall. The apparatus was controlled and the data were recorded using Med-PC IV computer software (Med Associates, Inc.).

### 3.3. Contexts

Two contexts were used that differed along visual, olfactory, and tactile dimensions. In one context, laminated sheets of black and white vertical stripes were mounted on the hinged front door and transparent wall of the chamber, a smooth black plexiglass sheet was positioned on the floor, and a paper towel with 1ml of 10% vanilla essence (Queen Fine Foods, Queensland, Australia) was placed in the bedding. In the second context, the hinged front door and wall were left clear, with a stainless-steel grid floor and a paper towel placed in the bedding, which had 1ml of 10% coconut essence (Queen Fine Foods, Queensland, Australia) added. Paper towels were changed prior to every session. The identities of Contexts A and B were fully counterbalanced across animals, such that Context A was the stripey-walled, smooth-floored, vanilla-scented context and Context B was the clear-walled, grid floor, coconut-scented context for half of the animals, and the remaining animals received the opposite arrangement.

### 3.4. Magazine training

Magazine training took place on day 1 in Context A. For these sessions, the house light was turned on at the start of the session and turned off when the session was terminated. No levers were extended. During the session, 20 deliveries of pellets and 20 deliveries of 20% sucrose solution were delivered on independent RT60 schedules, after which the session terminated.

### 3.5. Lever press training

Lever press training took place over 6 days (days 2-7) in Context A. Each session lasted for 50 minutes and consisted of two 10 minutes periods on each lever (i.e., four x 10 minutes sessions in total) separated by a 2.5 minutes time-out period in which the levers were retracted and the houselight switched off. Lever press periods terminated early if 20 outcomes were earned such that rats could earn a maximum of 40 pellets and 40 deliveries of sucrose solution per session. For half of the animals, the left lever earned pellets and the right lever earned sucrose, and the other half received the opposite arrangement (counterbalanced). For the first 2 days of lever press training, lever presses were continuously reinforced. Animals were shifted to a random ratio (RR-)5 schedule for the next 2 days (i.e. each lever earned an outcome with a probability of 0.2), then to a RR-10 schedule (i.e. each lever earned an outcome with a probability of 0.1) for the final 2 days.

### 3.6. Habituation

Rats were pre-exposed to Context B on day 8, following the last lever-press training session and prior to extinction training. This served to familiarize the animals to this context and reduce neophobia. Pre-exposure sessions lasted 40 minutes, during which no levers were extended and no food was delivered

### 3.7. Extinction

Rats were assigned to groups AAA, ABB, ABA and AAB after being matched for responding on the last day of acquisition training. For groups AAA and AAB, extinction training occurred in Context A. For groups ABA and ABB, extinction was conducted in Context B. Extinction sessions were 30 minutes long during which the houselight was turned on and both levers extended and lever presses recorded, but no outcomes were delivered.

Following termination of the extinction session, rats in Experiment 1 were put back into their home cage for 5 minutes to allow the experimenter to make any of the necessary context changes prior to reinstatement testing.

### 3.8. Outcome-selective reinstatement test

After 5 minutes in their home cages, rats were placed back into the operant chamber that was now adorned in the correct context for test. On test, both levers were available for the entire session and rats received 4 reinstatement trials separated by 4 minutes each. Each reinstatement trial consisted of a single free delivery of either the sucrose solution or the grain pellet. All rats received the same trial order: sucrose, pellet, pellet, sucrose. Responding was measured during the 2 minutes periods immediately before (Pre) and after (Post) each delivery. The reinstatement test session lasted for 30 minutes.

### 3.9. Tissue preparation and immunofluorescence

Two hours after the start of the reinstatement test, rats were deeply anesthetised via CO2 inhalation and perfused transcardially with cold 4% paraformaldehyde in 0.1 M phosphate buffer saline (PBS; pH 7.3-7.5). Brains were rapidly and carefully removed and postfixed in 4% paraformaldehyde overnight and then placed in a PBS solution containing 30% sucrose. Brains were sectioned coronally at 40 μm through the OFC, pDMS and DH defined by Paxinos and Watson (2014) using a cryostat (CM3050S, Leica Microsystems) maintained at approximately −20°Celsius. The sectioned slices were immediately immersed in cryoprotectant solution and stored at −20°Celsius. Five representative sections from each region of interest (mOFC, lOFC, pDMS and DH) were selected for each rat. Sections were first washed three times (10 minutes per wash) in PBS to remove any exogenous substances. The sections were then incubated in a blocking solution comprising of 3% Bovine Serum Albumin (BSA) + 0.25% TritonX-100 in 1x PBS for one hour to permeabilize tissue and block any non-specific binding. Sections were then incubated in anti-c-Fos primary antibody (1:500, Synaptic Systems Catalog #226 003) as an ‘activation marker’ (c -Fos is an immediate early gene that is expressed following neuronal activation) diluted in blocking solution for 72 h at 4°C. Sections were then washed 3 times in 1 × PBS and incubated overnight at 4°C in donkey anti -rabbit AlexaFluor-488-conjugated secondary antibody (1:1000, Invitrogen, Catalog #A21206). Every section was mounted on glass slides and were coverslipped using the mounting agent Vectashield and left to dry overnight in darkness. For quantification of c-Fos, a single image was taken of the mOFC/lOFC, pDMS, and DH CA1 per hemisphere of each slice (10 images in total per brain region of rat) on a Fluorescence Microscope (Zeiss) using a 10x air objective. Regions taken for each section were identical (2048 x 2048 pixels). Images were quantified using imaging software (ImageJ, Fiji Cell Counter). Briefly, background subtraction was applied to remove background noise. Images were then converted to binary, and thresholding was used to isolate stained cells. Finally, the Analyze Particles tool was used to quantify the number of cells based on a minimum particle size of 80 pixel units.

### 3.10. Data and Statistical analysis

Lever press and magazine entry data were collected automatically by Med-PC (version 5) and uploaded directly to Microsoft Excel using Med-PC to Excel software. Lever press acquisition and extinction data were analysed using repeated measures (Group x Session) ANOVA controlling the per-family error rate at α=0.05. For a more fine-grained analysis of test data, we used planned, complex orthogonal contrasts controlling the per-contrast error rate at α=0.05 according to the procedure described by Hays (Hays, 1973). If interactions were detected, follow-up simple effects analyses were calculated to determine the source of the interaction. Data were expressed as mean ± standard error of the mean (SEM) and averaged across counterbalanced conditions. Values of p < 0.05 were considered statistically significant. Statistical softwares SPSS and PSY were used to carry out these analyses. The data files and details of the statistical analyses for these experiments can be accessed at the following link: https://osf.io/52fv4/?view_only=137b51c2902245229864f8c2d84fff70.

## 4. Results – Experiment 1

### 4.1. Behavioural Results

Lever press acquisition is shown in Figure 2A, averaged across left and right levers. It is clear from this figure that all animals in Experiment 1 acquired the lever press response, and groups did not differ on their acquisition. This is supported by a main effect of day F(1,36) = 41.459, p = .00001, no main effect of group and no day x group interaction Fs < 1.

**Figure 2.**
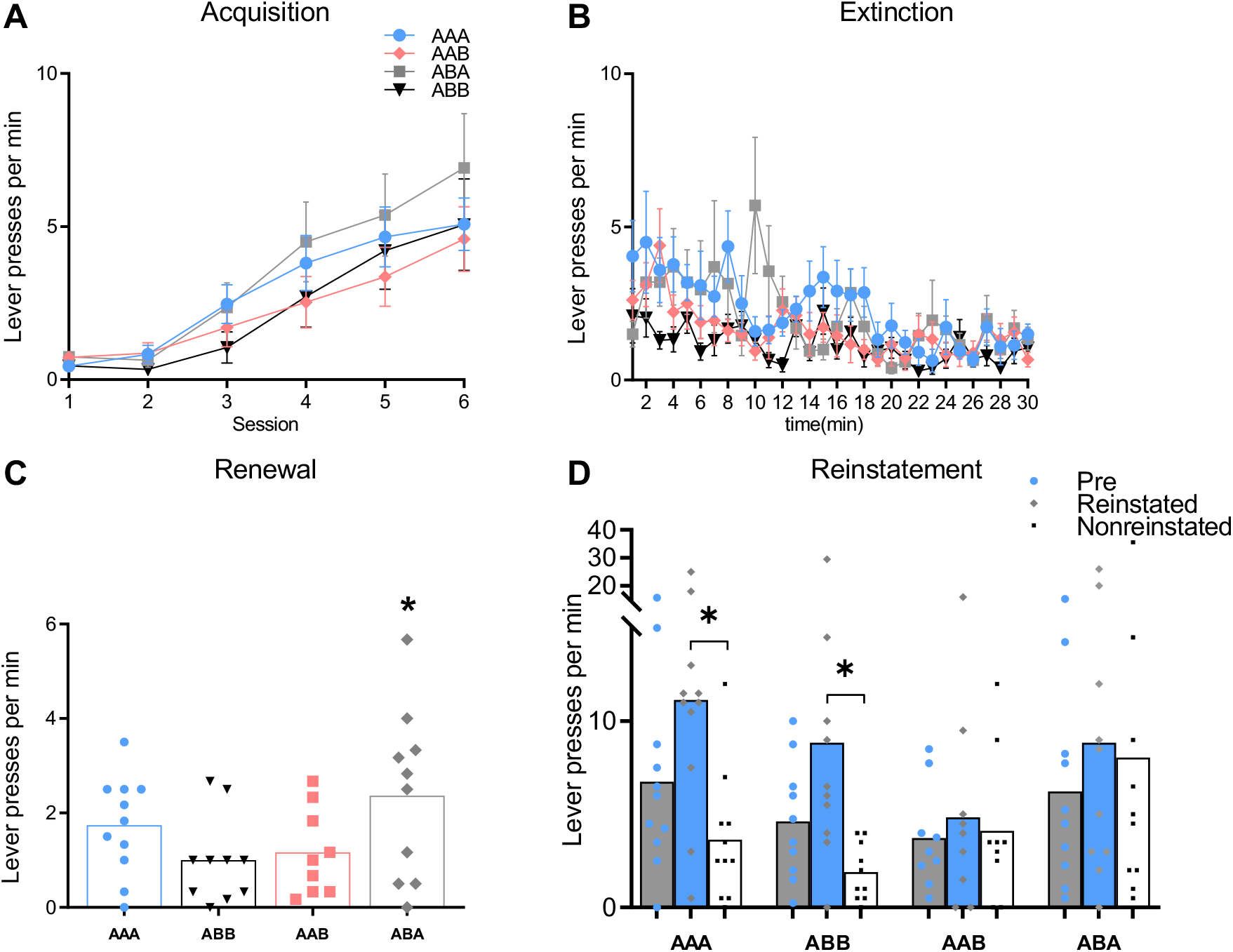
Lever presses per min (± SEM) during acquisition (A) and extinction (B). (C) Lever presses per min (± SEM) during first 3 minutes of the reinstatement test, prior to outcome delivery. (D) Lever presses per min (± SEM) during the reinstatement test. * p < .05

Responding during the 30 minutes extinction session is shown in Figure 2B. This figure shows that all animals reduced responding over this session, and that this did not differ between groups. This is supported by a main effect of minute (Greenhouse-Geisser corrected for violating sphericity), F(7.337,36) = 4.725, p = .000 that did not interact with group, F(22.012, 36) = 1.278, p = .185.

Responding during the first 3 minutes of reinstatement testing is shown in Figure 2C. Any increase in responding detected during this period can only be a result of context effects (i.e. ‘renewal’) and not reinstatement, because no outcomes were delivered during this period. Figure 2C shows that responding was highest in group ABA relative to the rest of the groups. Statistical analysis confirmed that this was the case (i.e. ABA > average [AAA/ABB/AAB]), F (1,36) = 5.596, p = .024. By contrast, responding in group AAB did not differ from groups AAA/ABB, F < 1, and groups AAA and ABB also did not significantly differ from each other, F(1,36) = 1.904, p = .176. These results suggest that there was an effect of renewal in group ABA but not in group AAB.

Performance during the reinstatement test is shown in Figure 2D. From this figure, it is clear that outcome-selective reinstatement was intact (Reinstated > NonReinstated) for groups AAA and ABB, and impaired (Reinstated = NonReinstated) for groups ABA and AAB. This is supported by a main effect of reinstatement F(1,36) = 10.213, p = .003, that did not interact with the AAA vs. ABB comparison, F < 1. Nevertheless, selective reinstatement was impaired for groups AAB and ABA, as demonstrated by a significant group (AAA/ABB vs AAB/ABA) x reinstatement interaction, F(1,36) = 6.691, p = .041. This interaction is supported by significant simple effects demonstrating greater responding on the reinstated than the nonreinstated lever in groups AAA, F(1,36) = 9.958, p = .003, and ABB, F(1,36) = 7.774, p = .008, but no evidence of differential responding on either lever in groups ABA and AAB, both Fs < 1.

### 4.2. Results of the c-Fos analysis

Representative photomicrographs demonstrating the extent of c-Fos expression in the medial OFC are shown in Figure 3A, in the lateral OFC are shown in Figure 3B, in the pDMS are shown in Figure 3C, and in the DH (roughly CA1 region) are shown in Figure 3D. Total c-Fos counts for these regions shown in Figures 3E (mOFC and lOFC), 3F (pDMS), and 3G (DH), respectively. Statistical analyses revealed c-Fos expression did not differ between groups in mOFC/lOFC or DH CA1 (p > 0.05). Our analysis did detect higher c-Fos expression in the pDMS for groups that demonstrated intact outcome-selective reinstatement on test (i.e. groups AAA and ABB) than for groups that did not (i.e. groups AAB and ABA), supported by a complex contrast (AAA/ABB vs AAB/ABA) demonstrating a main effect of group, F(1,36) = 32.394, p = .00001. Together, these results suggest that higher levels of neural activity in the pDMS, but not mOFC, lOFC, or DH, was associated with intact outcome-selective reinstatement.

**Figure 3.**
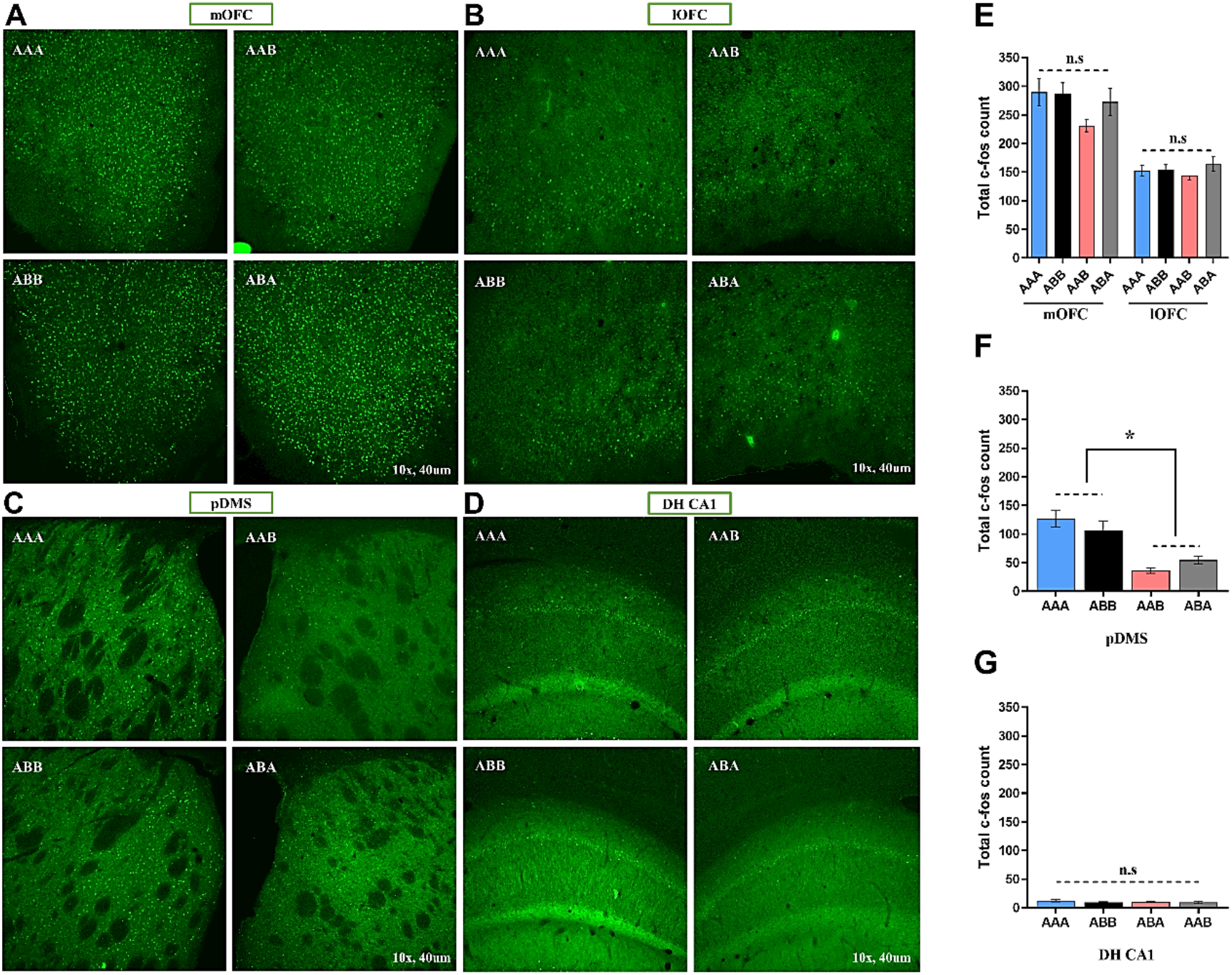
Representative immunofluorescence images of c-Fos levels in medial/lateral orbitofrontal cortex (A-B), posterior dorsomedial striatum (C), and dorsal hippocampus CA1 (D) for Experiment 1. E) Total c-fos counts (± SEM) for medial/lateral orbitofrontal cortex, F) Total c-fos counts (± SEM) for posterior dorsomedial striatum, G) Total c-fos counts (± SEM) for dorsal hippocampus CA1 region. * p < .05

## 5. Experiment 2: Outcome-selective reinstatement and extinction learning are both context-independent in an instrumental choice paradigm after multiple days of extinction training

In contrast to our expectations, the results of Experiment 1 suggest that outcome-selective reinstatement was only partially context-specific. This is because it was impaired in groups AAB and ABA but was intact in groups AAA and ABB, despite group ABB being tested in a context (B) in which outcome-response pairings had never been experienced. The major difference between groups for which selective reinstatement was intact (AAA and ABB) versus those for which it was impaired (AAB and ABA) was that extinction and testing occurred in the same context for the two former, but not the two latter, groups. Together, therefore, the results of Experiment 1 suggest that although the selective reinstatement effect appears to be independent of the context in which outcome-response associations are learned, the extinction learning upon which selective reinstatement relies is context-specific.

One potential problem with this conclusion, however, is the fact that we did not observe a renewal effect in group AAB. That is, if extinction learning was specific to the context in which it was learned, then it should not have transferred to Context B on test for this group such that renewal – consisting of elevated responding relative to groups AAA and ABB in the first 3 minutes of test (prior to outcome delivery) – should have been observed here, but it wasn’t. This failure to detect AAB renewal is notable for two reasons. First, as the current study is the first to our knowledge to investigate renewal in a paradigm involving instrumental choice, it could suggest that AAB renewal is not (but ABA renewal is) replicable in such a paradigm. Second, because selective reinstatement was impaired for group AAB despite the absence of a renewal effect, it could suggest that this impairment was independent of renewal rather than a consequence of it. It was the aim of Experiment 2 to investigate these questions.

An important difference between Experiment 1 and prior studies that have detected AAB renewal (Bouton & Ricker, 1994; Bouton et al., 2011; Tamai & Nakajima, 2000) is that we here conducted extinction and testing on the same day rather than on separate days. Although we chose this procedure based on the parameters we (Bradfield et al., 2015; Bradfield et al., 2018) and others (Ostlund & Balline, 2007) have used previously, almost all prior demonstrations of renewal and reinstatement have involved multiple days of extinction training, and testing on a separate day. It is possible that such a procedure could allow for better consolidation of the extinction context-no outcome association, resulting in more robust renewal effects on test. Thus, we decided to adopt these more traditional parameters in Experiment 2 in an attempt to increase the possibility of observing AAB renewal on test, and to determine if this led to impaired selectivity of reinstatement.

The design of Experiment 2 was identical to that of Experiment 1, except that rats were given 2 x 30 minutes extinction sessions over two days, and tested one day later. This time, we expected to observe a renewal effect in both groups ABA and AAB, as indicated by enhanced responding relative to groups AAA and ABB in the first 3 minutes of test. We further expected to replicate the findings that selective reinstatement was impaired for groups AAB and ABA relative to groups AAA and ABB.

## 6. Materials and Methods – Experiment 2

### 6.1. Animals

A total of 30 male and 30 female Long-Evans rats (n = 15 per group, N = 60) were used for Experiment 2. Male and female rats were distributed evenly between groups (n = 7 or 8 females, and n = 7 or 8 males per group). Animals were housed, food deprived, and maintained as described for Experiment 1. All procedures were approved by the Ethics Committees of the Garvan Institute of Medical Research, Sydney.

### 6.2. Apparatus and Contexts

All Apparatus and Contexts were as described for Experiment 1.

### 6.3. Behavioural Procedures

All behavioural procedures were conducted identically to that described for Experiment 1, except that rats were given 2 x 30 minutes extinction sessions for Experiment 2, which were conducted on different days (i.e. Days 9 and 10 of the experiment, following mag training, lever press training, and habituation to Context B). Reinstatement testing occurred one day later, on Day 11, identically to the manner described for Experiment 1.

### 6.4. Tissue preparation and immunofluorescence

All tissue preparation and immunofluorescence procedures were as described for Experiment 1.

## 7. Results – Experiment 2

### 7.1. Behavioural Results

Lever press acquisition is shown in Figure 4A, averaged across left and right levers. As is clear from this figure, all animals acquired the lever press response, and this did not differ by group. This is supported by a main effect of day F(1,56) = 108.711, p = .000, no main effect of group, F < 1) and no group x day interaction F < 1. Responding during the two 30 minutes extinction sessions is shown in Figures 4B-C. There was evidence of within-session extinction on both days, and this did not differ between groups. For the first extinction session, there was a main effect of minute (Greenhouse-Geisser corrected for violating sphericity), F(8.864,36) = 15.64, p = .00001 that did not interact with group, F(26.592, 56) = 1.163, p = .15 and a main effect of minute for the second session (Greenhouse-Geisser corrected for violating sphericity), F(11.996,56) = 3.862, p = .00001 that did not interact with group, F < 1. There was also a between-session extinction effect that did not differ by group, as evidenced by a main effect of day, F(1,56) = 100.276, p = .00001, and no group x day interaction, F < 1.

**Figure 4.**
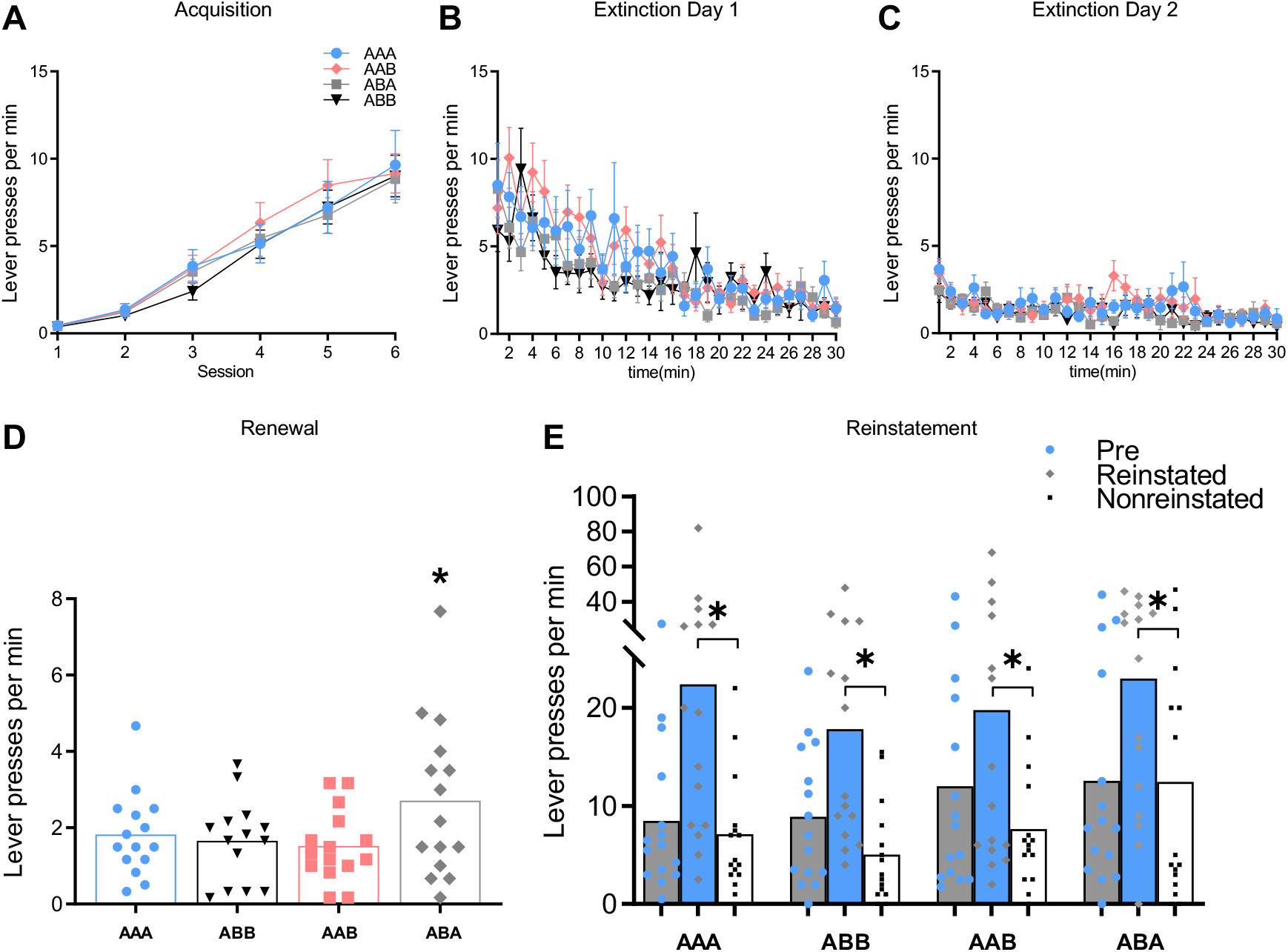
Lever presses per min (± SEM) during acquisition (A) and extinction (B-C). (D) Lever presses per min (± SEM) during the first 3 minutes of reinstatement testing, prior to outcome delivery. (E) Lever presses per min (± SEM) during the reinstatement test. * p < .05

Performance during the first 3 minutes of reinstatement testing is shown in Figure 4D. Note that data for one rat from group ABB was removed from this analysis due to fulfilling the outlier criteria of responding at a rate greater than three standard deviations above the mean. From this figure it is clear that, once again, there was a robust renewal effect in group ABA but not in group AAB. Statitiscally, group ABA again responded more than the other groups (i.e. ABA > average [AAA/ABB/AAB]), F (1,55) = 6.452, p = .016, whereas responding in group AAB did not differ from groups AAA/ABB, F < 1, nor between groups AAA and ABB, F < 1. This result indicates that we were once again unable to demonstrate AAB renewal in this multiple action, multiple outcome paradigm, even when we employed more traditional extinction/renewal parameters.

Performance during reinstatement testing is shown in Figure 4E. From this figure, and in further contrast to expectations, selective reinstatement was intact for all groups, including groups AAB and ABA that received extinction training and testing in different contexts. There was a main effect of selective reinstatement (Reinstated > Nonreinstated), F(1,56) = 35.94, p = .000, which did not interact with any group differences, all Fs < 1. We were so surprised by this result that we added n = 5 animals to each group (after initially replicating the group sizes of n = 10 from Experiment 1) to ensure that we were not underpowered to detect any small but significant effects between groups. However, the addition of these animals only strengthened our observations.

Together, these results demonstrate that outcome-selective reinstatement is entirely context-independent after multiple days of extinction training, and that AAB renewal is not replicable in a two action, two outcome, instrumental paradigm, at least with the parameters employed here. Moreover, the results of Experiment 2 appear to confirm that outcome-selective reinstatement and renewal effects are distinct phenomena. This is because in Experiment 1, selective reinstatement was impaired in group AAB despite the absence of a renewal effect, whereas in Experiment 2, selective reinstatement was intact in group ABA despite the presence of a renewal effect.

### 7.2. Results of the c-Fos analysis

As shown in Figure 5A-G, c-Fos expression did not differ between groups in any of the brain regions investigated for this experiment, all Fs < 1.

**Figure 5.**
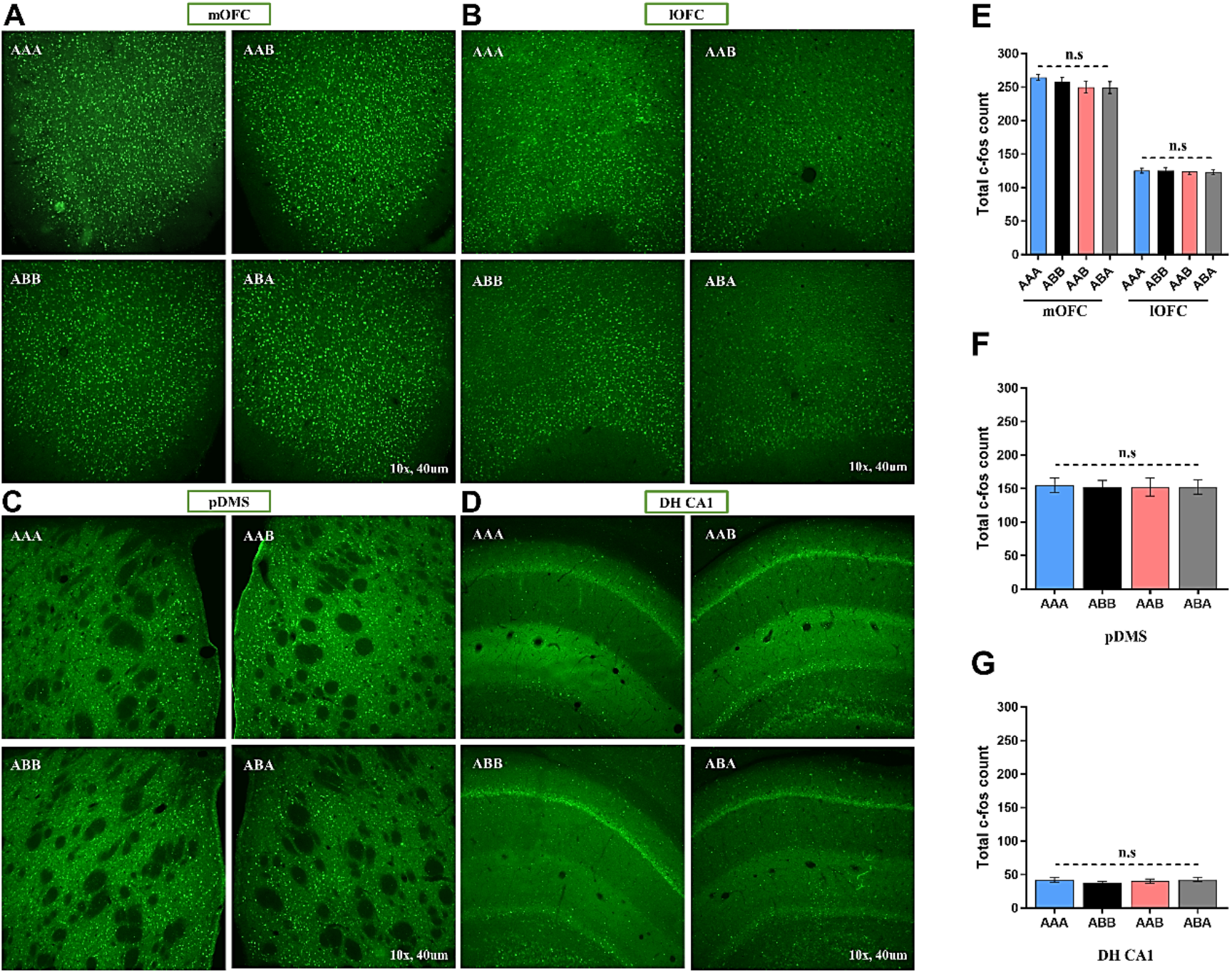
Representative photomicraphs of c-Fos levels in medial/lateral orbitofrontal cortex (A-B), posterior dorsomedial striatum (C), and dorsal hippocampus CA1 (D) for Experiment 2. E) Total c-fos counts (± SEM) for medial/lateral orbitofrontal cortex, F) Total c-fos counts (± SEM) for posterior dorsomedial striatum, G) Total c-fos counts (± SEM) for dorsal hippocampus CA1 region. * p < .05

## 8. Discussion

The experiments reported here reveal that the selective reinstatement of an action is partially context-independent, when tested immediately following extinction. By contrast, when tested after multiple days of extinction, outcome-selective reinstatement is entirely context-independent. Moreover, the context-specificity of reinstatement appears to be independent of increases in responding related to the context switch alone: - i.e. renewal. This is because impaired selectivity of reinstatement (i.e. Reinstated = Nonreinstated) was observed in group AAB in Experiment 1 despite the absence of a renewal effect in this group, whereas intact selectivity of reinstatement (i.e. Reinstated > Nonreinstated) was observed in Experiment 2 in group ABA, despite the presence of renewal. Finally, our results suggest that selective reinstatement is associated with neural activity in pDMS, but not the OFC or DH, because only in the pDMS did c-Fos expression reflect the intact/impaired nature of outcome-selective reinstatement.

Despite the unexpected nature of several of these results, one conclusion that can be confidently reached based on current findings is that the outcome-response associations that underlie outcome-selective reinstatement can be expressed independently of their physical context. That is, across both experiments, we observed selective reinstatement to be intact for group ABB despite the response and outcome never having been experienced in Context B prior to test. This was in contrast to our prior speculations made both here and elsewhere (Abiero & Bradfield, 2021), that selective reinstatement would be specific to the context in which outcome-response associations are initially learned, based on the fact that stimulus-response habits are generally context-specific (Bouton et al., 2011; Thrailkill & Bouton, 2015). This speculation relied upon the claim that outcome-response associations represent a special type of stimulus-response association, in which the outcome functions as a stimulus (Ostlund & Balleine, 2007). Current results appear to suggest, however, that outcome-response and stimulus-response associations are distinct from each other, at least with respect to their dependence on physical contexts. One possible reason for this distinction could be due to the nature of the ‘stimuli’ that enter into habitual stimulus -response associations relative to the nature of outcomes acting as stimuli. For example, lever-associated stimuli, such as the sight and smell of the lever, might be encoded as a part of the context in a way that outcomes are not. Indeed, levers in operant chambers are literally attached to the walls, whereas food outcomes can be lifted away from the food receptacle and consumed wherever the animal takes it. Thus, it is possible that outcome-response associations are encoded as being more distinct from their contexts, and thus might drive behaviour in a way that is distinguishable from (although still somewhat similar to) habits.

Our finding that selective reinstatement is predominantly context-independent does raise the question, however, as to why prior studies have found (non-selective) reinstatement of a single instrumental action to be diminished by alterations in context (Baker et al., 1991) when such reinstatement is presumably also underscored by outcome-response associations. One possibility is that the context might ‘gate’ the outcome-response association in a single action-outcome design, in a way that does not occur in a two action, two outcome design, although why this would be the case is unclear. Alternatively, it has been suggested (Adams, 1982; Holland, 2004) that training with a single action-outcome contingency results in the sensory properties of the outcome no longer controlling responding, which is controlled by its affective valence instead. By contrast, they suggested that training with two distinct action-outcome contingencies preserves the encoding of sensory properties. Thus, if we assume that affective valence is more likely to become attached to its context than the sensory properties of the outcomes are, then we would expect singular reinstatement to be more context-specific than selective reinstatement.

Another potential issue raised by the current results is why selective reinstatement was impaired for groups ABA and AAB in Experiment 1, but not in Experiment 2, when the only differences between these experiments was the amount of extinction training (1 vs. 2 sessions, respectively) and the length of time from the last extinction session to test (5 minutes vs. 24 hours, respectively). One seemingly straightforward way to interpret these findings would be to assume that although extinction learning did not transfer between contexts in animals tested immediately after brief extinction training, it did transfer in animals trained and tested over multiple days. Indeed, our own work provides some precedent to the idea that instrumental learning is initially context-dependent, but becomes context-independent over time (Bradfield et al., 2020), and further suggests that it is the additional time (as opposed to additional extinction training) between training and test that is the key variable in achieving such independence. Applying this interpretation to current results is complicated, however, by the fact that selective reinstatement and renewal effects appeared to be independent of each other, suggesting that the observation of selective reinstatement was not entirely dependent on the transfer of extinction learning. However, our ability to fully interpret each of these effects as reflective of *learning* is limited, as we can only base our conclusions on the observable *performanc*e of the animals, the latter of which is not a perfect indicator of the former. Moreover, it is possible that the change in experimental conditions from lever press acquisition, in which levers were trained separately, to extinction where both levers were presented simultaneously, provided an additional ‘contextual’ change that affected learning/performance, particularly in group AAB where renewal was expected but not observed. Future studies are therefore necessary to determine whether the contextual-impairment of selective reinstatement is an effect that is truly independent from renewal, and whether extinction learning is transiently context-dependent in instrumental choice learning as these results seem to suggest. Another area that requires further research, is the precise neural circuitry underlying the selective reinstatement effect. Unfortunately, our exploratory c-Fos analyses was not particularly illuminating with regards to this question, as we were only able to confirm the previously-demonstrated finding (Yin et al., 2005) that neural activity in the pDMS is related to selective reinstatement. We were not able to form any similar such conclusions for any of the other regions we studied, including medial and lateral OFC, and DH. The failure to find any differences in DH is particularly surprising, given the central role of this region in spatial and contextual representations. Once again, however, it is difficult to interpret a null effect, particularly in a correlative analysis such as this, so we cannot make any inferences about the cognitive mechanisms that might have led to (lack of) c-Fos expression in the DH in our experiments. Nevertheless, as very little is known about the neural circuitry of outcome-selective reinstatement, there is fertile ground for future studies to investigate this. We suggest that regions such as the basolateral amygdala and mediodorsal thalamus as excellent candidates for these studies, given their central roles in motivated behaviour.

Finally, we began this article by expressing a desire to contribute to the creation of a richer and more complex model of relapse than the single action, single outcome models currently available, so we will here conclude what our findings contribute to this understanding. Although we recognise that there are many steps that must be tackled before the work here can be thought to directly implicate certain features regarding humans with compulsive disorders, if we assume some degree of translatability that can be applied, then based on our findings we might predict that the likelihood of relapsing on two outcomes is higher when the abstinence has been brief and relapse occurs in a context other than the abstinence-associated context. Current results further imply that relapse should be more outcome-specific after longer periods of abstinence, regardless of context. To give the same example as before, if an individual has been abstinent from both smoking and drinking for a short time, and they relapse to smoking (or even breathe in passive smoke) as soon as they return to their local bar, they are also more likely to relapse on alcohol. If that individual has been abstinent for longer, however, their relapse is more likely to be specific, regardless of where they are.

## CRediT authorship contribution statement

Arvie Abiero: Conceptualization, Methodology, Investigation, Validation, Formal analysis, Writing – original draft

Zaid Ali: Investigation, Validation

Bryce Vissel: Project Administration, Funding Acquisition

Laura Bradfield: Conceptualization, Methodology, Formal Analysis, Supervision, Project Administration, Funding Acquisition, Writing – review and editing.

## Declaration of Competing Interest

None

## Acknowledgements

This work was supported by the Australian Research Council (DP200102445) awarded to L.A.B and B.V. We would like to thank Marios Panayi for helpful discussions regarding this manuscript.

## Notes

### Competing Interest Statement

The authors have declared no competing interest.

